# Finding *de novo* methylated DNA motifs

**DOI:** 10.1101/043810

**Authors:** Vu Ngo, Mengchi Wang, Wei Wang

## Abstract

Increasing evidence has shown that posttranslational modifications (PTMs) such as methylation and hydroxymethylation on cytosine would greatly impact the binding of transcription factors (TFs). However, there is a lack of motif finding algorithms with the function to search for motifs with PTMs. In this study, we expend on our previous motif finding pipeline Epigram to provide systematic *de novo* motif discovery and performance evaluation on methylated DNA motifs. Using the tool, we were able to identified methylated motifs in *Arabidopsis* DAP-seq data that were previously demonstrated to contain such motifs^1^. When applied to TF ChIP-seq and DNA methylome data in H1 and GM12878, our method successfully identified novel methylated motifs that can be recognized by the TFs or their co-factors. We also observed spacing constraint between the canonical motif of the TF and the newly discovered methylated motifs, which suggests operative recognition of these cis-elements by collaborative proteins.

## Introduction

DNA methylation is a major epigenetic mark that plays crucial roles in many key biological processes. Promoters with high DNA methylation levels are correlated with repressed genes, whereas promoters with low levels of methylation generally observe high levels of gene expression^2^. Previously, it was understood that DNA methylation correlates with low levels of TF-DNA binding and subsequently gene repression^2^. However, new studies have also shown that some TFs preferentially bind to methylated sequences and these sequences are often different from the canonical motifs of these TFs^3^. For example, Hu *et al* have found that Kruppel-like factor 4 (KLF4) can bind to CCmCGCC sequence (mC refers to the methylated cytosine), which is different from its canonical motif (CACACC)^3^.The protein RBP-J was also demonstrated to bind specifically to a methylated CpG-containing sequence^4^ (GmCGGGAA) in a methylation-dependent way. These observations illustrate the importance of identifying methylated motifs.

Currently, aside from Viner *et al.’s* work^5^, which was submitted at the same time to BioRxiv as our tool^6^, there is no other computational method to identify methylated motifs. Here we present a new version of our published motif finding method Epigram^7^ to identify methylated motifs and motifs containing other PTMs. Epigram can identify motifs in very large sets of sequences such as in a previous study in 980,465 sequences with a mean length of 1640bp^7^. Such a large set of sequences would be impractical for other motif finding programs to process. For example, HOMER would simply crash given such a data set^7,8^; MEME only accepts input size of ≤ 60000 characters with sequence lengths of ≤ 1000 base pairs^9,10^. Epigram’s scaling efficiency is comparable to that of DREME^11^ but it can find motifs longer than 8 base pairs, which is the default motif length in DREME^11^. The program discovers motifs by building position-specific weight matrices (PWMs) from the most enriched k-mers in the positive sequences over the negative sequences as “seeds” and extending the motifs to both directions^7^. We’ve expanded the alphabet to also represent PTMs such as methyl cytosine, carboxy cytosine, formyl cytosine, hydroxymethyl cytosine. This new algorithm provides a new tool that can identify DNA motifs with PTMs, which would be useful for analyzing the epigenomic data mapping different PTMs.

## Methods and Results

### Expansion on *de novo* motif discovery

The *de novo* discovery step requires input of both DNA sequences of interest and the PTM data such as DNA methylation data. It has two modes: one for finding motifs containing methylated CpG dinucleotides (mCpG), the other for other cytosine modifications such as non-CpG methyl cytosine and hydroxymethyl cytosine).

Since the vast majority of CpG methylations are symmetrical (i.e contain mC on both strands), the reverse compliment of a mCpG dinucleotide is itself. For example, the reserve compliment of ACGTmCG is mCGACGT. This feature allows us to conserve methylation information on a sequence’s reverse compliment without having to mark the guanine that pairs with the modified cytosine. In this mode, we add another base (E) to the conventional 4 bases (A, C, G, T) to represent mC, we call this mode *typeE.* In the second mode, we introduce another base, F, to mark the guanine that pairs with the modified cytosine *(typeEF).* This mode creates a more complex alphabet thus increasing the running time of the algorithm; therefore, we only allow for one type of modification at a time. The default seed length for this mode of mEpigram is 7 (instead of 8 for the methylated CpG mode).

Because of the introduction of the new bases, a new motif-scanning tool is needed in Epigram to search for matching k-mers using the modified motifs (m-motifs). To scan for the occurrences of a motif of interest in a set of DNA sequences, the program first simulates a score distribution for the motif by dinucleotide-shuffling the input sequences and calculates the scores for all of the k-mers inside the shuffled sequences using the motif’s PWM. The shuffling is repeated several times until an adequate number of k-mers is achieved (set to be 1 million in this study). Motif matches are called by passing a score threshold. This score threshold is defined by given p-values so that only a fraction equaling to the p-value can pass. For example, the score threshold for p-value of 0.01 is the lowest score in the top 1% of the k-mers from the shuffled sequences. The score of a k-mer given a motif (represented as a PWM) is calculated as:

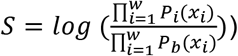

where *w* is the motif width, *P_i_*(*x_x_*) and *P_b_*(*x_i_*) are the probabilities of observing nucleotide *x_i_* at position *i* from the motif and the background distributions, respectively.

### Performance evaluation with simulation

To evaluate the performance of mEpigram in retrieving positive signals, we conducted several rounds of simulation tests. We first chose the ChIP-seq peak regions of a TF EGR1 in the H1 cells (8743 sequences at 260bps per sequence on average). We then inserted methylated motifs in these sequences and check how many of them can be correctly retrieved. We generated the inserted k-mers from a PWM and we varied the information content (IC) of the PWM from 0.23 to 2 per position to check the capability of mEpigram to retrieve strong and weak modified motifs (**Table 1**). Furthermore, we also tested the algorithm on various motif’s enrichment levels by inserting k-mers into 5%, 2%, 1% and 0.5% of randomly selected regions and try to retrieve the motif with mEpigram. To calculate the similarity between a pair of the original retrieved m-motif and the retrieved one, we aligned them by sliding one PWM over another and calculated the Pearson correlation coefficient for all the alignments with as least 4bp overlap. The maximum Pearson correlation coefficient represents the motif similarity. In our test, a pair of motifs is considered the same if the correlation is at least 0.95.

**Table 1:** Retrieval of inserted motifs in the simulated data. Abundant or highly conserved motifs can be identified more easily compared to ones that are scarce and have low information content.

| Motif | IC per position | % of peaks |  |  |  |
| --- | --- | --- | --- | --- | --- |
|  |  | 0.5% | 1% | 2% | 5% |
| Motif 1 | 2.0 |  |  |  |  |
| Motif 2 | 1.05 | Not found |  |  |  |
| Motif 3 | 0.63 | Not found | Not found |  |  |
| Motif 4 | 0.38 | Not found | Not found |  |  |
| Motif 5 | 0.23 | Not found | Not found | Not found | Not found |

The enrichment of a motif is measured by the ratio of number of peaks in the input regions that contain the motif of interest over the number of peaks in the shuffled regions that contain it. To determine a significant enrichment level, we generated a distribution of possible enrichment scores using shuffled sets of sequences. The distribution is approximately normal; the enrichment score of 1.5 is approximately 3 standard deviations from the mean (**Figure S.1**). Therefore, we chose 1.5 as the enrichment cutoff for our motifs.

Depending on the IC of the motifs, mEpigram can identify m-motifs that are present in as low as 0.5% of the sequences (in the case of completely conserved motif with IC of 2) or not able to find motif even at 5% abundance (in the case of least conserved motifs with IC of 0.23) (**Table 1**). Since most known motifs have average ICs of about 1.0^12^ (standard deviation of 0.035), we expect the program to be able to find motifs with abundances as low as 1%.

### Identifying m-motifs in known methylation-prefer TFs

O’Malley et al. have demonstrated certain TFs do prefer methylated sequences when bind with DNA. We obtained 270 TF DAP-Seq datasets from this study to test our algorithm’s ability to find m-motifs in validated data. The TFs were categorized based on their sensitivity toward changes in methylation levels. This was calculated as the change in binding affinity after removing methylation from the sequences^1^. Out of 270 TFs, 11 are considered to bind to methylated sequences preferentially, 66 have no preference between methylated and non-methylated DNA, 193 are disrupted by methylation.

We used TF binding specificity to simulate TF binding affinity. The binding specificity of a TF motif is defined as the ratio of the density of motif matches within the TF-binding regions over the density of motif matches found in the genome, excluding the TF-binding regions. The importance of methylation in a motif is measured by the logarithm of the fold change in binding specificity before and after removing methylation information from an m-motif (logFC). A significant m-motif is thus defined as: 1) appears in at least twice as many regions in the input sequences compared to the dinucleotide-shuffled sequences. 2) when methylation information is removed, m-motif’s binding specificity is reduced by at least 2.83 fold, which corresponds to logFC of 1.5. Since 5-methylcytosine (5mC) in plants, as opposed to mammals, also occur largely in CHG context^13^, *typeEF* of the algorithm is more appropriate as it doesn’t assume only CpG methylations.

As the results, we were able to find significant m-motifs for all of the 11 TFs reported to bind preferentially to methylated DNA. Out of 66 TFs that were considered insensitive to DNA methylation, 7 of them contain significant m-motifs. Only 1 out of 193 TFs known to be disrupted by DNA methylation have m-motifs. Without this additional filter, the numbers are respectively: 11, 40, and 46.

### Application of mEpigram to TF ChIP-seq data

#### Retrieval of m-motifs

It is widely accepted that DNA methylation disrupts binding of TFs but recently studies suggested that some TFs may prefer methylated motifs (e.g CEBPB^14^). mEpigram provides a tool to study the impact of DNA methylation on TF using the in vivo ChIP-seq binding data. To this end, we applied mEpigram to 55 TF ChIP-seq data in H1 and 44 datasets in GM12878 generated by ENCODE together with the whole genome bisulfite sequencing data. In the mEpigram runs, the maximum number of motifs to be reported was set at the default 200. Motifs from the same run were aligned to each other and redundant motifs were removed. Since the data we use only contains CpG methylation information, we take advantage of the *typeE* mode as it can handle longer k-mers, thus offers higher sensitivity.

First of all, mEpigram successfully found the canonical motifs in majority of the experiments, which indicates the success of the motif-finding algorithm. In H1, thirty-five out of 40 known canonical motifs were correctly identified by mEpigram in the top 5 most enriched motifs from the corresponding TF ChIP-seq experiments (**Table S2.1**). For GM12878, 24 out of 31 known canonical motifs are identified (**Table S2.2**).

The number of m-motifs found for each TF ranges from 0 to 41 (**Table S2.2**). Out of 55 ChIP-seq datasets in H1, 31 show enrichment for m-motifs >1.5 fold (**Table S2.2**). For GM12878, 24 out of 44 datasets have significantly enriched m-motifs.

The presence of m-motif can have different meanings for each TF. A TF can preferentially bind to methylated sequences or simple tolerate methylations. For example, the top m-motif motif of CEBPB in H1 is highly enriched and very similar to that of the canonical motif. It is identified at the majority of the canonical motif’s loci. For NRF1, the top m-motif does appear at the canonical motif’s loci but less frequently (**Figure S2.1**). There is a sharp increase in methylation level at the center of the motif-matching loci (**Figure S3**). This suggests that the canonical CEBPB motif binds preferably to methylated sequences. However, the methylation levels around the matching loci for CEBPB’s canonical motif is significantly lower than that of its m-motif, which shows CEBPB does not require methylation to bind with its motif **Figure S2.2**. For NRF1, the canonical motif prefers regions of lower methylation levels, whereas the m-motif is found in regions with higher methylation level (**Figure S3**). In contrast to CEBPB, there is no spike in the center of the plot for NRF1’s canonical motif. It can be interpreted that the NRF1 TF does not prefer methylated sequences, but it is insensitive to methylation.

For m-motifs that are different from their TF’s canonical motifs, for example CTCF and EGR1, they do not cooccur in the same ChIP-seq peaks with their canonical motifs often (**Figure S2.3-4**). CTCF and EGR1’s m-motifs are present in regions with high level of methylation (about 0.8) while their canonical motifs prefer low methylation levels (0.1-0.2, **Figure S3**). Given that the ChIP-seq peaks tend to occur in low methylation levels, the fact that these peaks have high methylation levels suggests that these m-motifs act as a way to counter-act the DNA methylation, thus recruit the TF. We hypothesize that removing the methylation of these m-motifs will disrupt the TF’s binding to these loci.

In most of these cases, the m-motif is significantly different from the canonical one or the most enriched motif present and some examples are shown in **Table 2**.

**Table 2:**
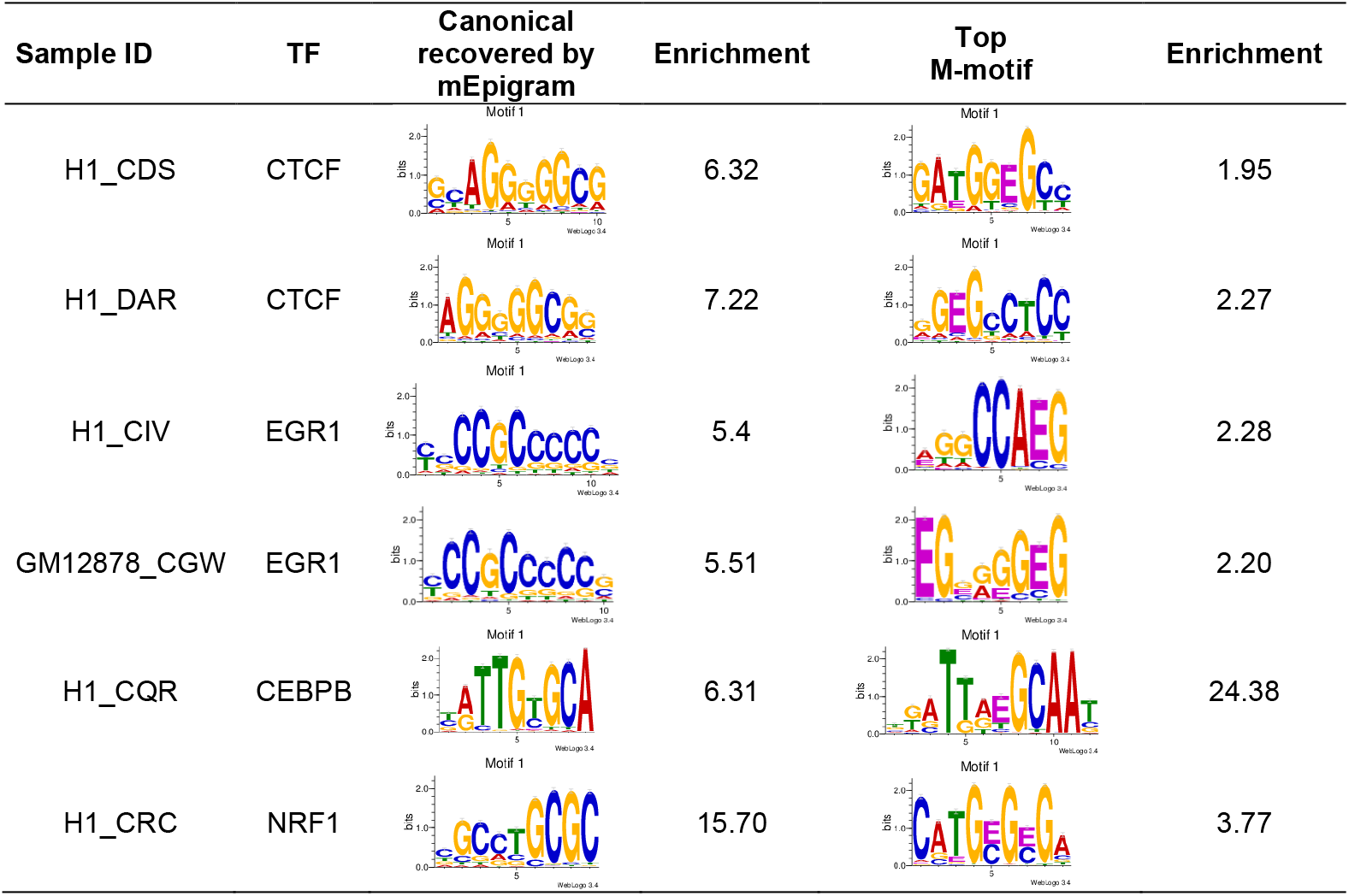
Datasets with significant m-motifs: Number of motifs and m-motifs of some of the samples with the most enriched m-motifs. See Supplemental table S2.2 for the complete list.

| Sample ID | TF | Canonical recovered by mEpigram | Enrichment | Top M-motif | Enrichment |
| --- | --- | --- | --- | --- | --- |
| H1_CDS | CTCF |  | 6.32 |  | 1.95 |
| H1_DAR | CTCF |  | 7.22 |  | 2.27 |
| H1_CIV | EGR1 |  | 5.4 |  | 2.28 |
| GM12878_CGW | EGR1 |  | 5.51 |  | 2.20 |
| H1_CQR | CEBPB |  | 6.31 |  | 24.38 |
| H1_CRC | NRF1 |  | 15.70 |  | 3.77 |

Some of the TFs in **Table 2** have been previously shown to either have interactions with DNA methylation or bind with specific methylated DNA sequences. For instance, CTCF is known to bind to DNA in a methylation specific manner and CTCF binding is regulated partly by differential DNA methylation^15^. The top m-motifs for two replicates of CTCF in H1 are similar to each other (Pearson correlation is 0.931 when aligned together with several bp shift). The third CTCF experiment in H1 (H1_CIU) used a different antibody and a different m-motif was found.

For CEBPB, the top motif is the methylated canonical CEBPB motif, which is consistent with the previous observation that CEBPB binds to 39% of the methylated canonical sequence^14^. We also discovered a strong m-motif for NRF1, present in 3.68% of the peaks, that is the canonical motif methylated at its CG dinucleotide. As a comparison, the canonical motif is present in 25% of the sequences. This finding is consistent with the observation that NRF1 TF exhibits binding with methylated sequences^3^.

#### Importance of cytosine

Furthermore, we evaluated the importance of cytosine methylation in the identified m-motifs. For each sample, we first de-methylated the m-motifs identified by mEpigram. In each PWM, at each position *i,* the probability of *E, P_i_* (*x_i_* = *E*), was added to *P_i_*(*x_i_ = C*) and then *P_i_* (*x_i_* = *E*) was set to zero. The resulted PWMs were next scanned against their respective peak regions without the methylation information (containing only A, C, G, T). Some of the m-motifs, when scanned after de-methylation, retain their enrichment (**Table 5**). This is often because these m-motifs are the methylated canonical motifs. For example, CEBPB and NRF1’s top m-motifs are both methylated canonical motifs. Their enrichments remain relatively unchanged after de-methylation. This further suggests that DNA methylation doesn’t hinder the TFs bindings. In contrast, some m-motifs have their enrichment significantly reduced after de-methylation (**Table 5**). These motifs generally contain more than 1 methylated cytosine in their sequences. Thus, removing the methylation significantly changes their enrichment. These motifs are likely selectively bound by their TFs.

**Table 5:**
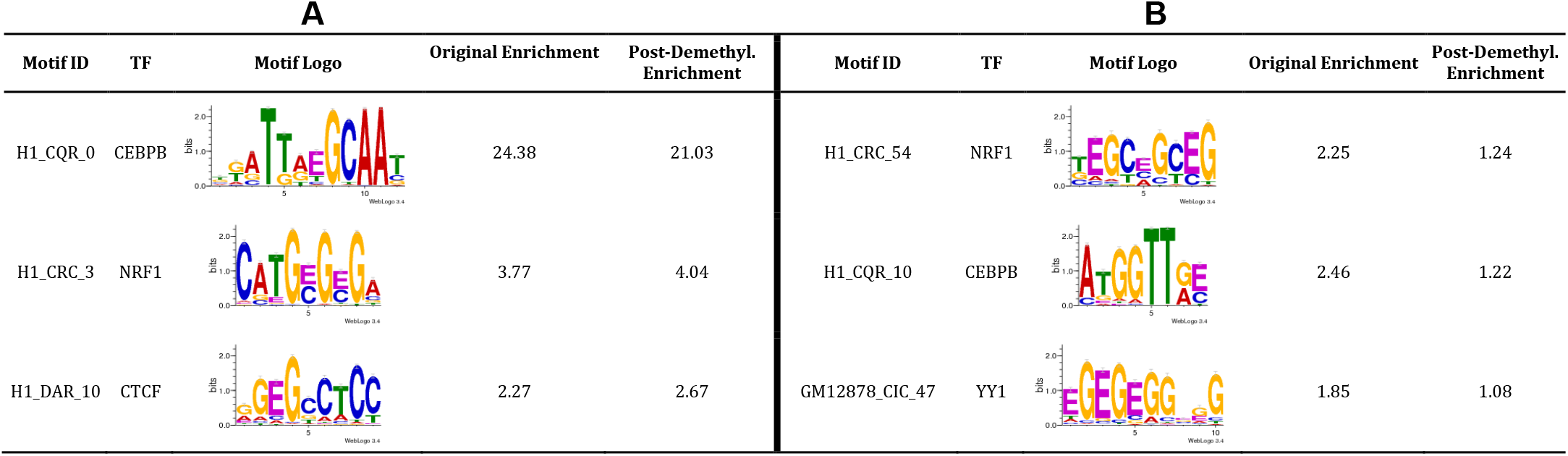
**A**/ Examples of the m-motifs retaining high enrichments after de-methylation. **B**/ Examples of the m-motifs enriched in their methylated form but not after de-methylation.

| A |  |  |  |  | B |  |  |  |  |
| --- | --- | --- | --- | --- | --- | --- | --- | --- | --- |
| Motif ID | TF | Motif Logo | Original Enrichment | Post-Demethyl. Enrichment | Motif ID | TF | Motif Logo | Original Enrichment | Post-Demethyl. Enrichment |
| H1_CQR_0 | CEBPB |  | 24.38 | 21.03 | H1_CRC_54 | NRF1 |  | 2.25 | 1.24 |
| H1_CRC_3 | NRF1 |  | 3.77 | 4.04 | H1_CQR_10 | CEBPB |  | 2.46 | 1.22 |
| H1_DAR_10 | CTCF |  | 2.27 | 2.67 | GM12878_CIC_47 | YY1 |  | 1.85 | 1.08 |

#### Distance constraints

We next searched for distance constraints between methylated motifs and other motifs using the SpaMo algorithm^16^ (**Table 3**). RepeatMasker^17^ was used to mask repeated sequences with chains of “N” to reduce false positives. Some m-motifs exhibit highly significant spacing constraints with other motifs, most notably the motifs identified from CTCF binding peaks. These CTCF motifs have enrichments over 1.5 and the SpaMo analyses gave the adjusted p-values of less than 0.001.

**Table 3:**
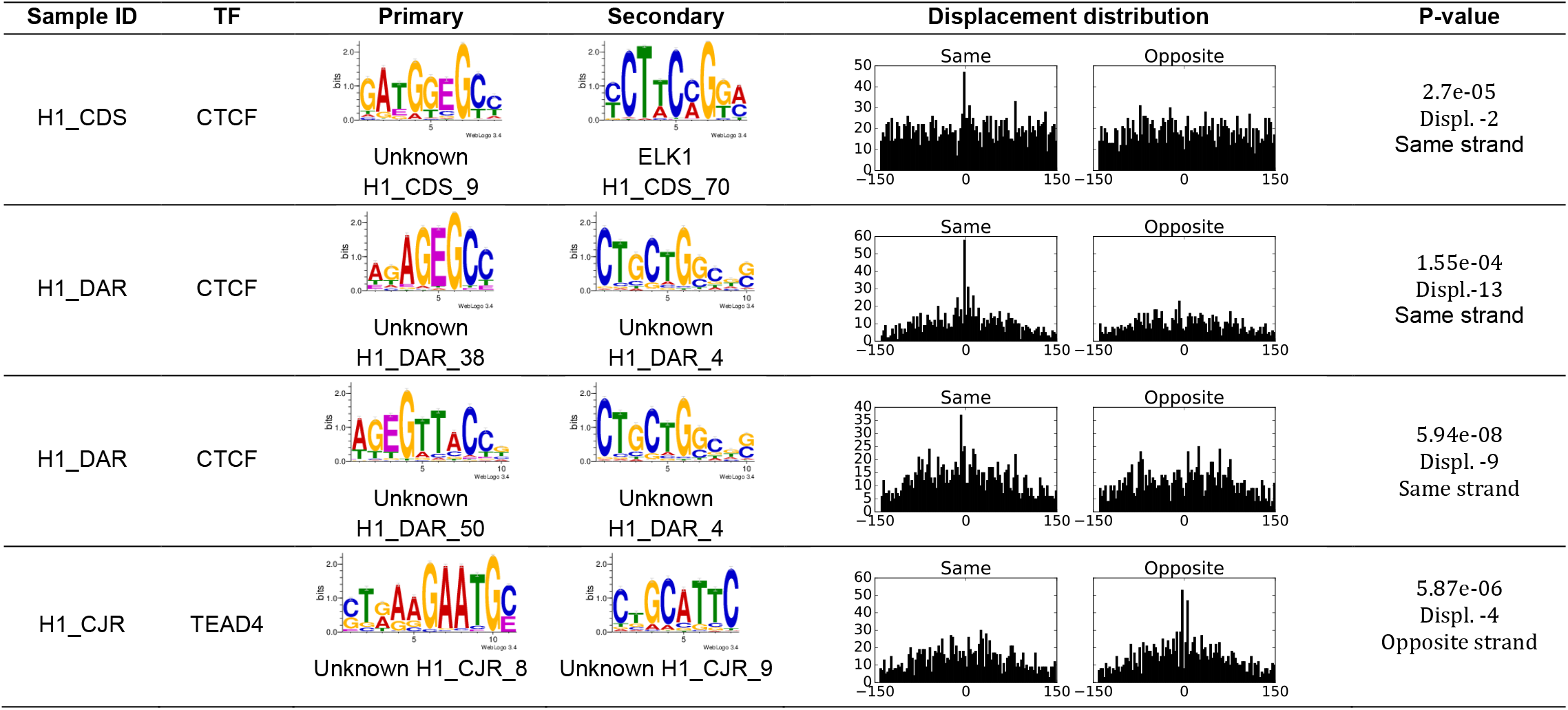
Most significant spacing pairs between m-motifs and other motifs. The motifs were scanned against Tomtom’s database and only matches with E-value less than 0.1 were accepted. The distance is the base pairs between the 3’ end of the primary motif and the 5’ end of the secondary motif. The p-value cutoff for significant displacement was set at 0.001. The histograms show the distributions of displacements between the primary and secondary motifs. The X-axis is the displacement, the Y-axis is frequency.

| Sample ID | TF | Primary | Secondary | Displacement distribution | P-value |
| --- | --- | --- | --- | --- | --- |
| H1_CDS | CTCF | Unknown<br>H1_CDS_9 | ELK1<br>H1_CDS_70 |  | 2.7e-05<br>Displ. -2<br>Same strand |
| H1_DAR | CTCF | Unknown<br>H1_DAR_38 | Unknown<br>H1_DAR_4 |  | 1.55e-04<br>Displ. -13<br>Same strand |
| H1_DAR | CTCF | Unknown<br>H1_DAR_50 | Unknown<br>H1_DAR_4 |  | 5.94e-08<br>Displ. -9<br>Same strand |
| H1_CJR | TEAD4 | Unknown H1_CJR_8 | Unknown H1_CJR_9 |  | 5.87e-06<br>Displ. -4<br>Opposite strand |

### Comparison of m-motifs in H1 and GM12878

To identify m-motifs that are common or unique in H1 and GM12878, we scanned motifs found in H1 peaks against GM12878 peaks and vice versa. The enrichments of the m-motifs in the other cell type were calculated and compared with the enrichment in the cell type where they were discovered. For CTCF, ERG1, and NRF1, several m-motifs are enriched in both GM12878 and H1. These motifs have enrichments of over 1.5 and appear in at least 2% of the peaks in both of the datasets. The top NRF1 m-motifs found in H1 and GM12878 are very similar to each other (**Table 4**). The top m-motifs for EGR1, on the other hand, are different from one another. The top CEBPB m-motifs found in in H1 were enriched in GM12878. In general, the motifs found in GM12878 for CEBPB appear significantly less often, the maximum enrichment is 4.7 compared to 24.38 in H1 (**Table 4**). This is unsurprising given the differences between the H1 and GM12878 data.

**Table 4:**
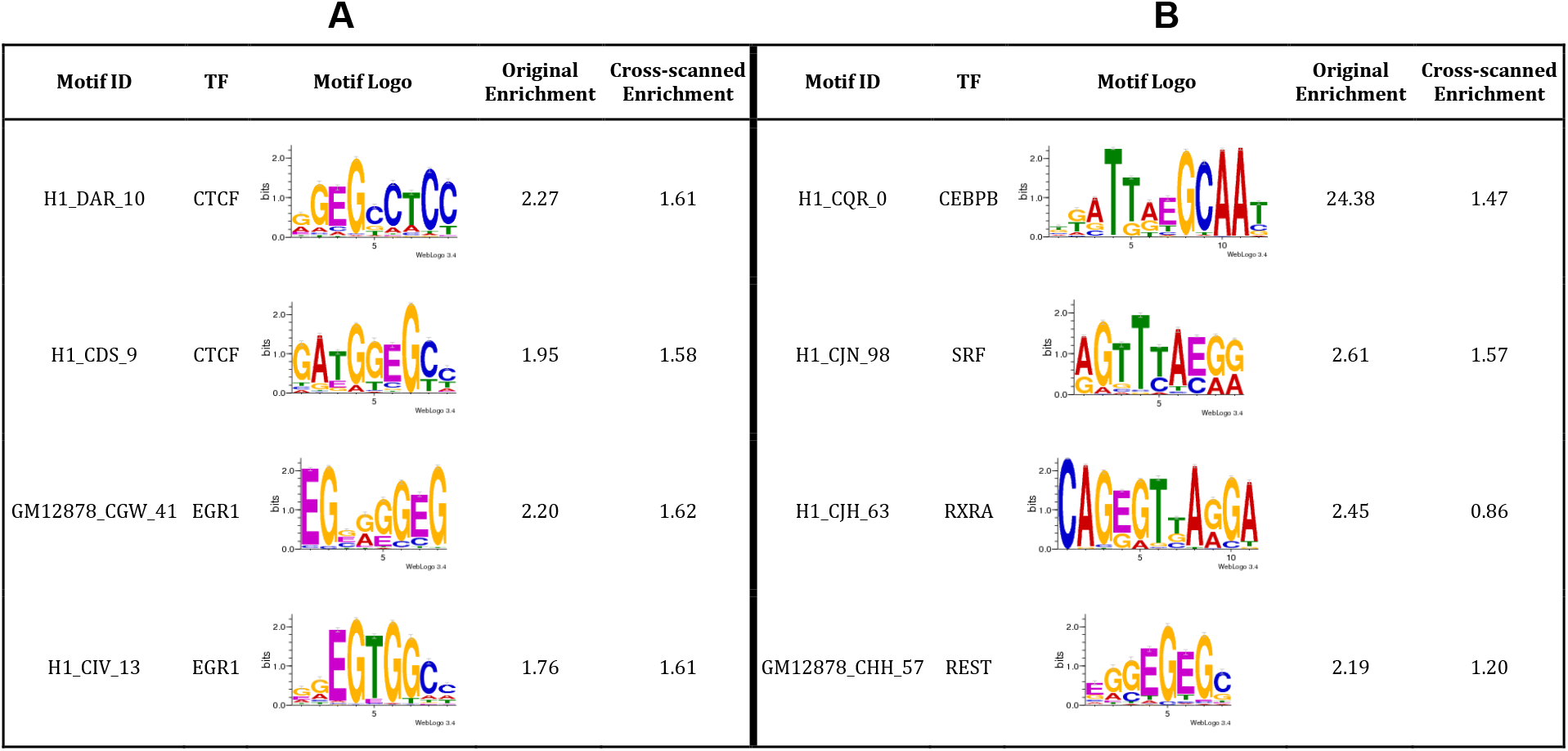
**A**/ Some of the consistently enriched m-motifs between H1 and GM12878: The enrichments remain significant when scanned in both H1 and GM12878 data. **B**/ Some of the differentially enriched m-motifs between H1 and GM12878.

| A |  |  |  |  | B |  |  |  |  |
| --- | --- | --- | --- | --- | --- | --- | --- | --- | --- |
| Motif ID | TF | Motif Logo | Original Enrichment | Cross-scanned Enrichment | Motif ID | TF | Motif Logo | Original Enrichment | Cross-scanned Enrichment |
| H1_DAR_10 | CTCF |  | 2.27 | 1.61 | H1_CQR_0 | CEBPB |  | 24.38 | 1.47 |
| H1_CDS_9 | CTCF |  | 1.95 | 1.58 | H1_CJN_98 | SRF |  | 2.61 | 1.57 |
| GM12878_CGW_41 | EGR1 |  | 2.20 | 1.62 | H1_CJH_63 | RXRA |  | 2.45 | 0.86 |
| H1_CIV_13 | EGR1 |  | 1.76 | 1.61 | GM12878_CHH_57 | REST |  | 2.19 | 1.20 |

### Effect of 5hmC on mEpigram results

Because 5-hydroxymethylcytosine and 5-methylcytosine are undistinguishable by bisulfite sequencing, we examined the effect of 5hmC’s presence on our findings. Using TAB-seq data for H1^18^, we found that the large majority of CG loci only contain 5hmC in low probability (95% CG loci having less than 0.1, with the mean of 0. 045). Removing 5hmC from H1 methylation data did not significantly change the mEpigram results.

## Discussion

We present here one of the first attempts to expand alphabet in motif search to meet the need of integrative analysis of sequence and epigenomic data. We have demonstrated the power and usefulness of mEpigram in identifying methylated motifs when combining sequence and DNA methylation. mEpigram can readily consider other modifications such as 5hmC, 5fC and 5cacC. When applied to analyzing human TF ChIP-seq data, mEpigram found that several TFs have significantly enriched methylated motifs. The most enriched m-motifs are the methylated canonical motifs (CEBPB, NRF1), which suggests that these TFs may be tolerant or prefer binding to their methylated sequences. Furthermore, additional novel m-motifs, that are not necessarily as enriched as canonical motifs, were also found in many TF binding regions (CTCF, EGR1). Particularly interesting, some of these methylated motifs are significantly enriched in the methylated form compared to the unmethylated form, which suggests possible impact of TF binding by methylation.

The effect of 5hmC on the bisulfites-sequencing data is not significant enough to produce false positives on mEpigram results. Therefore, bisulfite-sequencing data is adequate to identify m-motifs.

Author contributions. Vu Ngo modified the Epigram program, performed the analyses, and wrote the manuscript, Mengchi Wang contributed to the data analysis and manuscript preparation, Wei Wang conceived the idea, designed the project, interpreted the data and wrote the manuscript.

**Figure 1:**
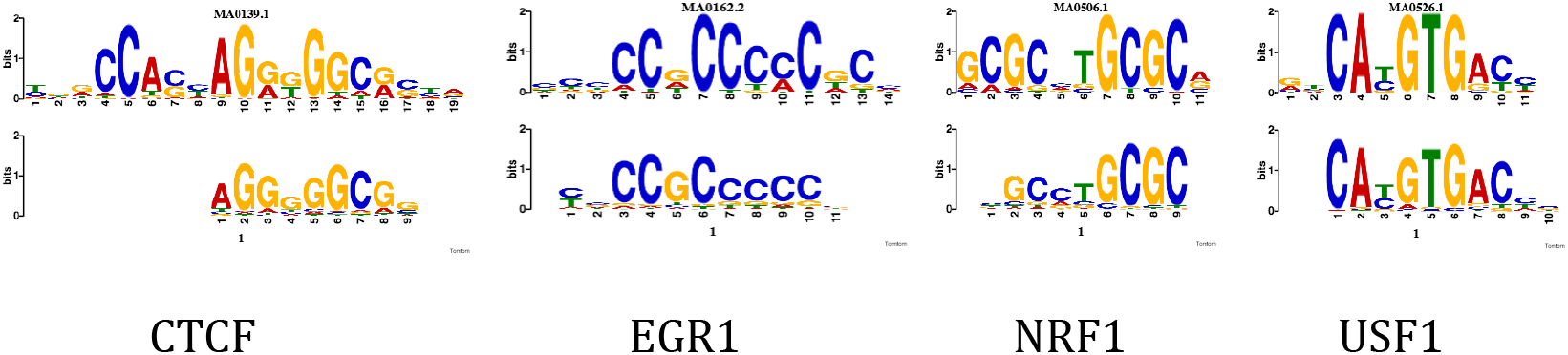
Top ranking non-methylated motifs found by mEpigram are consistent with canonical motifs. For each alignment, the top image is the canonical motif; the bottom image is mEpigram’s result. The top motifs found by mEpigram are compared to other databases using TomTom. Matches with p-value lower than 10e-6 are reported.

## Acknowledgement

This work is partially supported by NIH (U54HG006997) and CIRM (RB5-07012).

